# Human immunodeficiency virus (HIV) dynamics in secondary lymphoid tissues and the evolution of cytotoxic T lymphocyte (CTL) escape mutants

**DOI:** 10.1101/2023.03.10.532137

**Authors:** Wen-Jian Chung, Elizabeth Connick, Dominik Wodarz

## Abstract

In the secondary lymphoid tissues, human immunodeficiency virus (HIV) can replicate both in the follicular and the extrafollicular compartments. Yet, virus is concentrated in the follicular compartment in the absence of antiretroviral therapy, in part due to the lack of cytotoxic T lymphocyte (CTL)-mediated activity there. CTL home to the extrafollicular compartment, where they can suppress virus load to relatively low levels. We use mathematical models to show that this compartmentalization can explain seemingly counterintuitive observations. First, it can explain the observed constancy of the viral decline slope during antiviral therapy irrespective of the presence of CTL in SIV-infected macaques, under the assumption that CTL-mediated lysis significantly contributes to virus suppression. Second, it can account for the relatively long times it takes for CTL escape mutants to emerge during chronic infection even if CTL-mediated lysis is responsible for virus suppression. The reason is the heterogeneity in CTL activity, and the consequent heterogeneity in selection pressure between the follicular and extrafollicular compartments. Hence, to understand HIV dynamics more thoroughly, this analysis highlights the importance of measuring virus populations separately in the extrafollicular and follicular compartments rather than using virus load in peripheral blood as an observable; this hides the heterogeneity between compartments that might be responsible for the particular patters seen in the dynamics and evolution of the HIV *in vivo*.

## Introduction

The majority of HIV replication occurs in the secondary lymphoid tissues, such as lymph nodes, spleen, or the gut-associated lymphoid tissue (GALT) [1]. Within these tissues, virus replication is concentrated in the B cell follicles, with lower levels of replication observed in the extra-follicular compartments in the absence of antiretroviral therapy [2-13]. The follicular compartments contain follicular CD4 T helper cells with higher permissiveness for viral replication than extrafollicular CD4 T cells, and have a high concentration of extracellular virions on the surface of follicular dendritic cells (FDCs). Both aspects promote viral spread. Another reason for the higher concentration of virus in the B cell follicles, however, is that these compartments are relatively immune privileged from CTL [4,5]. CTL concentrate in the extrafollicular compartment, to which CTL home and in which they are stimulated. Therefore, virus load is lower in the extrafollicular compartment, controlled by CTL, while virus load is significantly higher in the follicular compartments, where CTL are substantially less abundant. This has been underlined with experiments in SIV-infected macaques, where the discrepancy in virus load between the two compartments is most pronounced in animals that show strong CTL-mediated control [4,5,14], and where the distribution of virus across the two compartments is more equal in animals with weaker CTL responses or during the acute phase of infection, before CTL responses have fully matured [4,5,14].

This unequal degree of CTL-mediated virus control in the follicular and extrafollicular compartments has implications for the evolutionary dynamics of CTL escape mutants. It is thought that CTL escape contributes to the loss of virus control over time as the disease progresses [15]. Point mutations occur in CTL epitopes, such that they are not recognized anymore by the responding CTL clone. A number of studies have pointed out that while CTL escape can occur rapidly during the initial stages of infection, the emergence of CTL escape occurs at a surprisingly slow rate in the longer term during chronic infection [16-20]. Several reasons have been suggested to explain this phenomenon. In mathematical models, a broad CTL response directed against multiple epitopes of the virus, or a fitness cost associated with escape, can render the evolution of escape more difficult to achieve, leading to a longer time until escape is observed [21-23]. Further, mathematical models suggest that the latent viral reservoir can slow down evolutionary processes, and this might also contribute to a delayed emergence of CTL escape if the selective advantage of the escape mutant is not too high [24]. Another line of reasoning has been that anti-viral CTL responses in HIV infection are weaker than previously thought. While CTL responses have been shown to significantly suppress virus load, it is possible that these responses act predominantly in a non-lytic way, suppressing viral infection or replication rather than killing infected cells [19,25]. Mathematical analysis of escape evolution data showed that the relatively slow rate of escape emergence can be consistent with that notion [19,25]. This has been further underlined by studies showing that the rate of SIV decline during anti-viral therapy was identical in in CTL-competent and CTL-depleted macaques [26,27], arguing against the role of CTL in shortening the life-span of infected cells. Similarly, the notion that CTL act largely in a non-lytic rather than a lytic manner was also suggested as an explanation for the observed constancy of the estimated infected cell life-span during anti-viral therapy in people living with HIV, which was found to be around 1-2 days across different patients regardless of disease stage/ CD4 T cell count [28].

Here, we use mathematical models to show that the unequal distribution of CTL activity in the follicular and extrafollicular compartments of the secondary lymphoid tissues can significantly impact virus dynamics and the rate of CTL escape evolution in HIV-infected patients. First, compartmentalization can explain the observed constancy of the estimated infected cell life span during antiretroviral therapy, because estimates are based on peripheral blood measurements that are likely influenced most by virus in the follicular compartments, due to the high viral loads there. Second, compartmentalization can lead to significantly slower rates of CTL escape emergence compared to dynamics where overall virus load is the same but cells and viruses mix extensively. The reason is as follows. While CTL-mediated selection pressure on the virus is strong in the extrafollicular compartment, virus load is low, which leads to a low chance for the virus to generate new escape mutants. In the follicular compartment, on the other hand, the chances for the virus to generate escape mutants are high, due to the large amount of virus replication that occurs there. Lack of significant CTL activity in this compartment, however, results in the lack of a strong selection pressure in this location, which in turn results in a low chance for escape mutants to grow and to migrate to the extrafollicular compartment.

### Basic model of compartmental dynamics

We use a mathematical model that describes the dynamics between HIV and CTL responses in the follicular and extrafollicular compartments [29]. The population of uninfected cells, infected cells, and CTL in the extrafollicular compartment are denoted by X_e_, Y_e_, and Z_e_, respectively. The corresponding populations in the follicular compartment are denoted by X_f_, Y_f_, and Z_f_, respectively. The time evolution of these populations is given by the following set of ordinary differential equations.

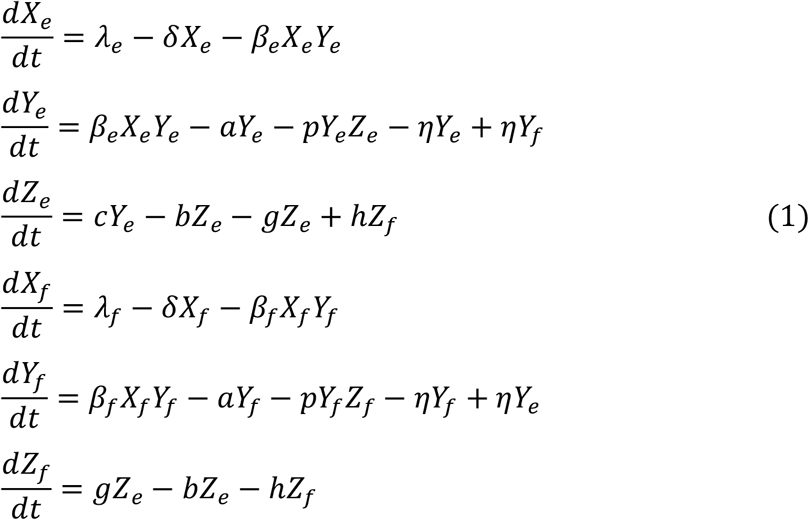

In each compartment, uninfected cells are produced with a rate *λ*_*e*_(*λ*_*f*_). These cells die naturally with a rate *δX*_*e*_ (*δX*_*f*_), and become infected with a rate *β*_*e*_*X*_*e*_*Y*_*e*_ (*β*_*f*_*X*_*f*_*Y*_*f*_). Infected cells die with a rate *aY*_*e*_ (*aY*_*f*_), and are killed by CTL with a rate *pY*_*e*_*Z*_*e*_ (*pY*_*f*_*Z*_*f*_). Infected cells migrate to the other compartment with a rate *ηY*_*e*_ (*ηY*_*f*_). CTL become stimulated and expand in the extrafollicular compartment with a rate *cY*_*e*_. The rate of expansion is assumed to be not proportional to the number of CTL, since this has been shown to result in more stable and hence more realistic dynamics [30]. The CTL migrate from the extrafollicular to the follicular compartment with a rate *gZ*_*e*_. In the follicular compartment, no CTL stimulation or expansion is assumed to occur, and CTL migrate back into the extrafollicular compartment with a rate *hZ*_*f*_. In both compartments, CTL are assumed to die with a rate *bZ*_*e*_ (*bZ*_*f*_). To achieve the observed scenario that most CTL are located in the extrafollicular compartment, we assume *g >> h*.

### Basic model properties

This model has been analyzed in some detail in [29]. For the limiting case *η* → 0, we can define the basic reproductive ratio of the virus in each compartment as R_0e_ = λ_e_β_e_/*δ*a, and R_0f_ = λ_f_β_f_/*δ*a, respectively. If both are larger than one, the infection is established in both compartments. In the absence of immunity (Z_e_=Z_f_=0), the following equilibrium is attained.

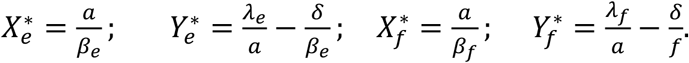

For *η* > 0, an infection in both compartments is established if

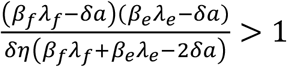

The equilibrium expressions in this case are too complicated to write down, but converge to the above expressions for low values of *η*, which is a biologically realistic assumption. If the population of CTL is added, they will expand as long as infected cells are present in the extrafollicular compartment, with the extent of the CTL expansion depending on the infected cell population size. The system then converges towards a stable equilibrium describing CTL-mediated virus control, which is again too complicated to write down. As with other CTL models [31], equilibrium virus load is inversely proportional to the strength of the CTL response, which in this model is determined by the rate of CTL expansion or CTL responsiveness (parameter *c*), and the rate of CTL-mediated killing (parameter *p*).

### Lytic CTL activity and the constancy of the infected cell life-span during ART

Our model only takes into account lytic CTL-mediated activity and ignores non-lytic suppression of virus replication by CTL. The reason is that we would like to investigate the dynamics under lytic activity, for which it has been more difficult to explain the evolutionary dynamics of the virus. Previous models of HIV dynamics with a lytic CTL response differentiated between two stages of infected cells, a cellular eclipse phase, and a virus production phase that is subject to CTL-mediated killing [22]. This allowed the model to reproduce the observed behavior that the overall decline rate of infected cells during anti-viral therapy remains relatively constant despite variations in the strength of CTL-mediated lysis. In our model, this distinction is not required to account for the constancy in the rate of virus decline in the face of variations in the CTL response. It arises naturally from the uneven distribution of CTL in the extrafollicular and follicular compartments. Figure 1 uses our model to simulate the initial dynamics of infected cell decline during anti-viral therapy (modeled by setting all infection rates to zero), brought about by the death of productively infected CD4 T cells. The infected cell decline is shown both in the presence and the absence of a CTL response. In the EF compartment, a clear difference is seen: The initial rate of virus decline is faster in the presence compared to the absence of CTL (Figure 1A), due to the effect of CTL-mediated killing on the life-span of infected cells. Over time, the rate of virus decline in the presence of CTL slows down, due to a reduction of the CTL population during treatment, when antigenic stimulation diminishes (not shown). In the F compartment, on the other hand, the rate of infected cell decline in the presence and absence of CTL are indistinguishable, due to lack of significant CTL-mediated activity in this location (Figure 1B). Because virus load is much higher in the F compared to the EF compartment (due to lack of CTL activity), the rate at which the total amount of virus (summed over both compartments) declines is dictated by the dynamics in F. In other words, the rate of decline of overall virus load is very similar both in the presence and absence of the CTL response (Figure 1C). Virus load measured in the peripheral blood likely reflects the sum of virus from the F and EF compartments. Hence, our model can account for the constancy of the viral decline slope irrespective of the strength of the CTL response, even though it assumes that CTL-mediated killing significantly reduces virus load (through action in the EF compartment). Note that we do not simulate the longer term virus decline dynamics during anti-viral therapy, since this would require a more complex model that tracks infected macrophages and latently infected cells [32]; this is not within the current scope of the analysis.

**Figure 1.**
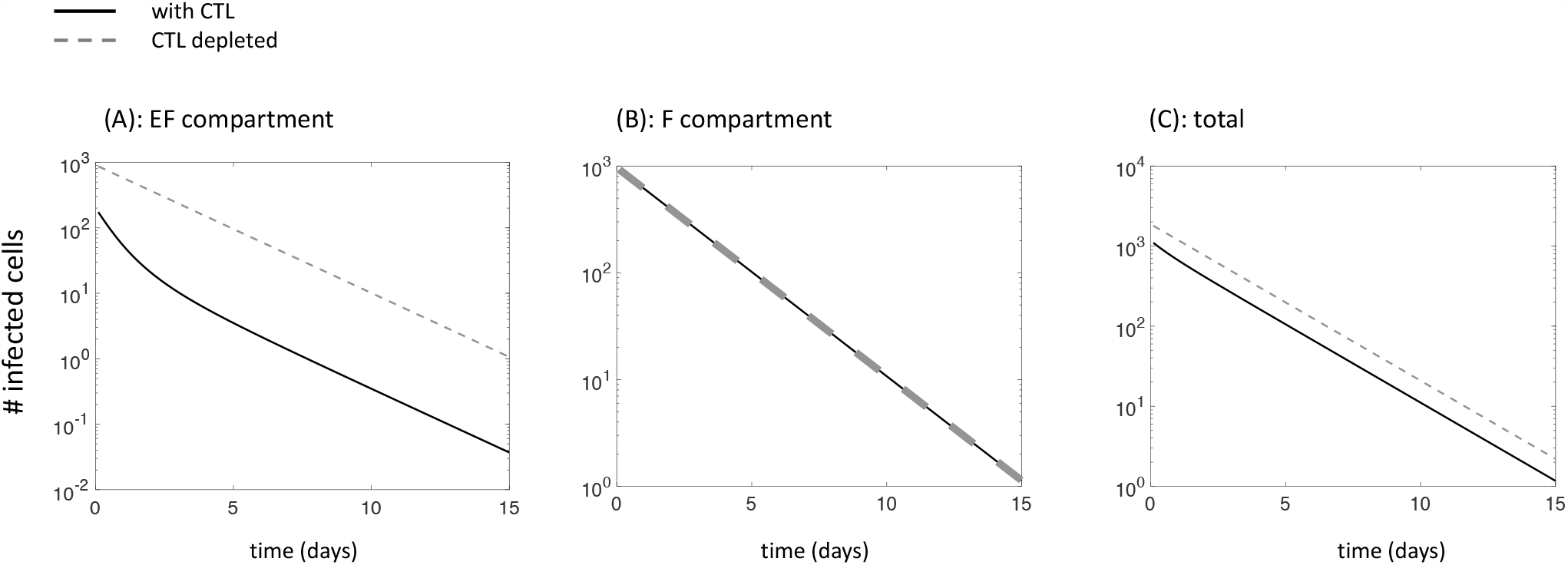
Simulation of ODE model (1) during antiviral therapy (*β*_*e*_*=β*_*f*_*=0*), in the presence (solid line) and absence (dashed line) of CTL. (A) Dynamics in the extrafollicular compartment. (B) Dynamics in the follicular compartment. (C) Total dynamics, where populations are summed up over both compartments. Parameters were chosen as follows. *λ*_*f*_*=500*; *λ*_*e*_*=500*; *δ=0.01*; *a=0.45*; *β*_*e*_*=β*_*f*_*=7×10*^*-5*^; *p=0.05*; *b=1*; *c=0.1*; *η=0.0001*; *g=0.0001*.

### Modeling evolution of CTL escape

Here, we introduce a population of mutant virus, which is not recognized by CTL anymore. Again, this virus can replicate both in the extrafollicular and the follicular compartments, and the corresponding infected cells are denoted by *Y*_*1e*_ and *Y*_*1f*_, respectively. The escape variant is generated by mutation during wild-type virus replication with a rate µ, and replicates itself with a rate *β*_*1e*_*X*_*e*_*Y*_*1e*_ (*β*_*1f*_*X*_*f*_*Y*_*1f*_). The resulting ordinary differential equations are given as follows.

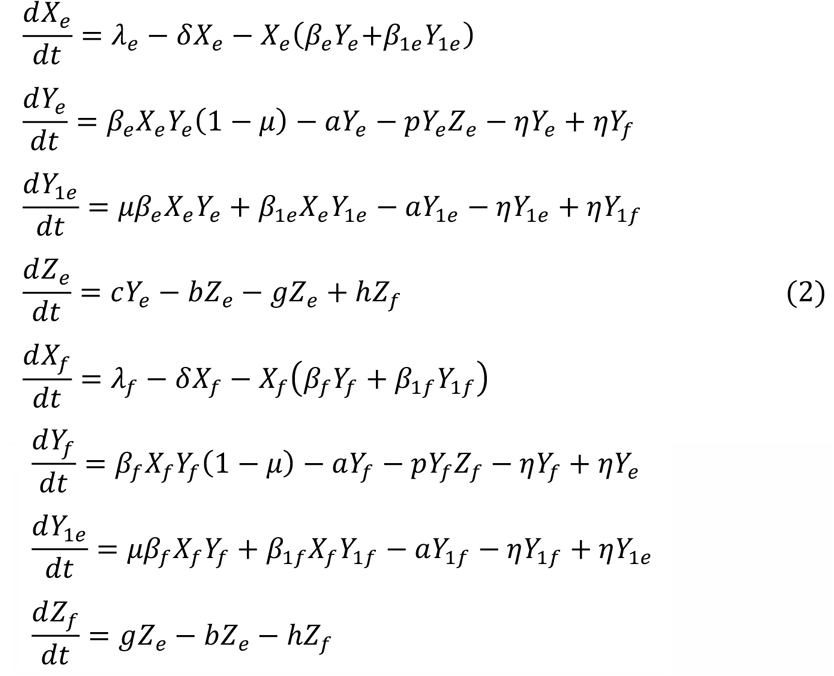

CTL escape mutants can carry a fitness cost compared to the wild-type virus in the absence of CTL [15], which is typically manifested in the parameter *β*. These mutants, however, tend to acquire compensatory mutations [15] that greatly reduce this fitness cost or eliminate it. We do not model this process here, because it is not essential to the question under investigation, and because this simplification limits the complexity of the model. We will hence assume a relatively small fitness cost of the escape mutant, expressed in a slightly lower value of *β*_1e_ (*β*_1f_) compared to *β*_e_ (*β*_f_).

### Basic outcomes of the model

The outcomes of this model are determined by the overall fitness of the escape mutant in the two compartments. Let us first consider a single compartment (see Supplementary Materials), i.e., the EF compartment where CTL become stimulated and kill (in the model, this would correspond to *η=0* and *g=0*). If the fitness cost of the escape mutant lies below a threshold, such that *β*_1e_ > *β*_e_(1-µ), then the escape mutant will fixate, i.e. it persists while the wild-type virus population goes extinct. The required fitness cost is very small and means that the escape mutant has to be almost neutral with respect to the wild-type virus (because µ is of the order 2×10_-5_[33]). If this condition is violated, on the other hand, both the escape mutant and the wild-type will coexist at a level determined by the strength of the CTL response. If the CTL response is relatively strong the escape mutant becomes the dominant virus strain since it is advantageous in this case. This dominance of the mutant can come close to fixation. If the CTL response is weak, however, the escape mutant will form only a small fraction of the infected cell population because its overall fitness is lower than that of the wild-type virus. It is essentially maintained by selection-mutation balance. For the two-compartment model, we assume a fitness cost of CTL escape mutants such that *β*_1e_ < *β*_e_(1-µ) such that complete mutant fixation does not occur. First consider strong compartmentalization, i.e. a low rate of virus exchange, *η*, between compartments and a low rate of CTL migration, *g*, into the F compartment (Figure 2A). This is a biologically realistic regime. In the presence of a strong CTL response that suppresses virus in the EF compartment, we see that the escape mutant dominates in the EF compartment, while it remains a minority in the follicles due to the lack of CTL activity and hence selection in that location (Figure 2Ai). This can correspond to the early chronic phase of infection. If the strength of the CTL response lies below a threshold, the escape mutant remains a minority in both compartments (Figure 2Aii). This might apply to advanced disease where immunity is already significantly weakened.

**Figure 2.**
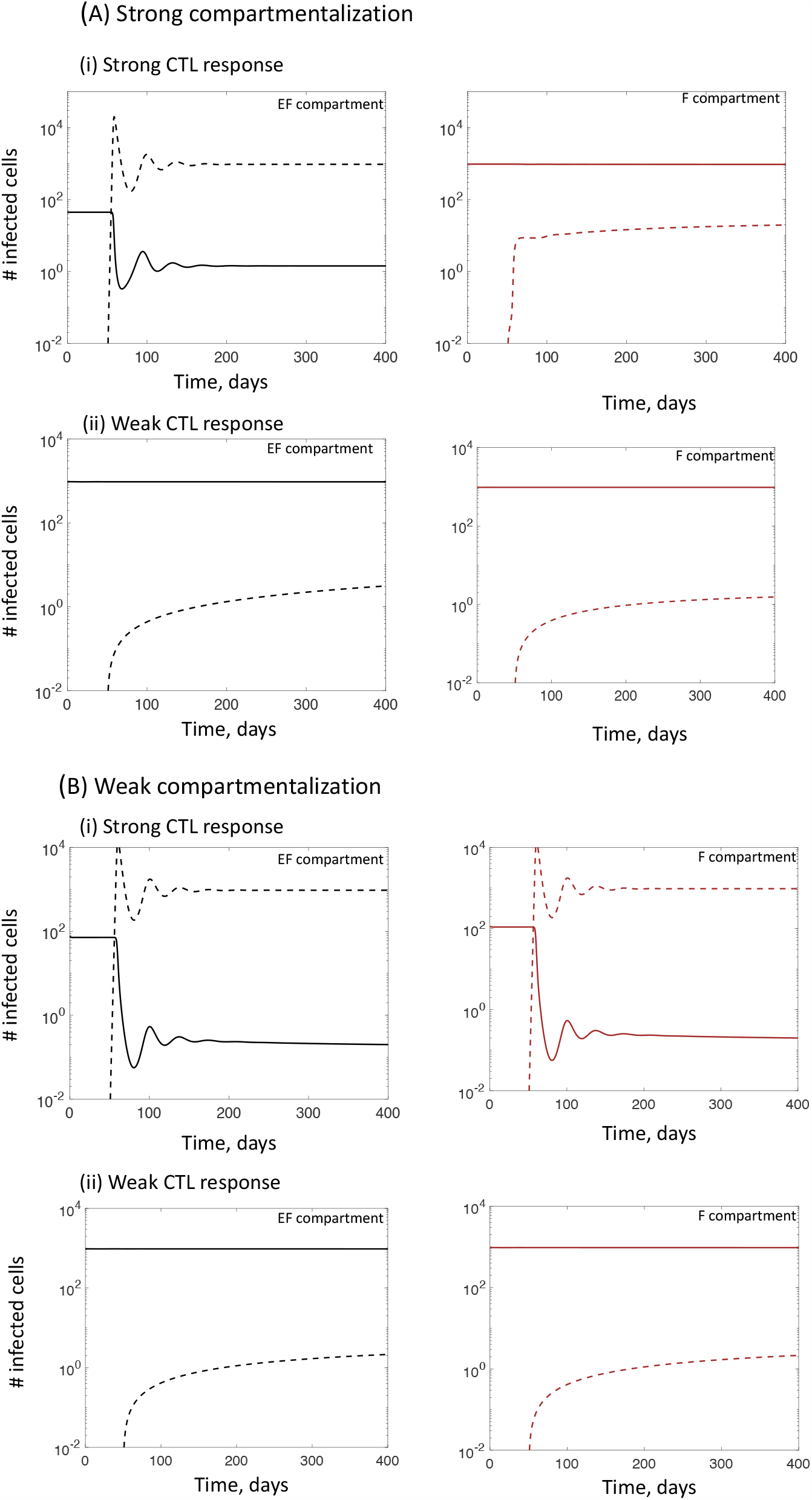
Outcomes in ODE model (2), including a CTL escape mutant. (A) These plots assume strong compartmentalization, i.e. infected cells move between compartments with a relatively slow rate, and CTL enter the follicular compartment with the slow rate. Dynamics are shown for (i) a relatively strong CTL response and (ii) for a relatively weak CTL response, determined by the parameter *c*. Left and right graphs show dynamics in the extrafollicular and follicular compartments, respectively. Solid lines depict the dynamics of the dynamics of the wild-type virus, dashed lines the dynamics of the CTL escape mutant. (B) Same graphs, but under the assumption that infected cells move with a faster rate between compartments and CTL move to the follicular compartment with a high rate, identical to the rate at which they move from the F to the EF compartment. Parameters were chosen as follows. *λ*_*f*_*=500*; *λ*_*e*_*=500*; *δ=0.01*; *a=0.45*; *β*_*e*_*=β*_*f*_*=7×10*^*-5*^; *β*_*1e*_*=β*_*1f*_*=0.99β*_e_; *µ=2×10*^*-5*^; *p=0.05*; *b=1*. For (A) *η=0.0001*; *g=0.0001*. For (B) *η=1*; *g=1*. For strong CTL responses, *c=1*. For weak CTL response *c=10*^*-4*^.

The strong compartmentalization can be compared to a highly mixed system where virus and CTL migrate with a relatively fast rate between compartments (larger values of *η* and *g*, see Figure 2B). This is not a biologically realistic scenario, but serves as a point of comparison. In this case, the outcome in both compartments is always identical because the CTL are more or less equally distributed. If the CTL response is strong (high value of c), the escape mutant dominates in both compartments (Figure 2Bi). For relatively low values of c, the escape strain represents only a minority in both compartments (Figure 2Bii).

### Stochastic simulations

So far, we have discussed the properties of the ODEs. Because the mutants initially exist at low numbers, however, stochastic effects should be taken into account. Therefore, we performed stochastic Gillespie simulations [34] of the model. As initial conditions, only the wild-type virus was assumed to be present, with populations being at equilibrium (they fluctuate around equilibrium values due to stochasticity). Over time, de novo mutations are produced. Eventually, for our parameter values, the escape mutant will rise to domination in the extrafollicular compartment because the CTL-mediated selection pressure there. We determined the average time until the CTL escape mutant reaches 95% of the infected cell population in this location, based on repeated realizations of the computer simulation. Only mutant domination in the EF compartment is considered, because this will contribute most to an increased overall virus load (since in the F compartment, it is assumed that the virus is not well controlled by CTL in the first place).

### Parameter values and assumptions

Parameter values of the simulations are given in the figure legends. The life-span of infected cells was set to be around 2 days [35], and the remaining parameters were adjusted such that the basic reproductive ratio of the virus, R_0_, is around 8 [36], and such that populations persist robustly in the stochastic Gillespie simulations. The cell population sizes in follicles and extralfollicular compartments are not well known, but on a qualitative level, the results presented here do not depend on these numbers. The mutation rate of the virus was assumed to be 2×10^−5^ per bp per generation, consistent with the order of magnitude observed experimentally for HIV [33]. In the base model, we assume that the rate of CTL migration into the F compartment, as well as the rate of infected cell exchange between compartments is low, such that we have a strongly compartmentalized model structure (parameters *g* and *η* are low), consistent with biological reality [4]. We will refer to this as **model version (i)**. For this model version, we explore how properties, particularly the average time to mutant invasion, depend on the strength of the CTL response, expressed by the parameter c, i.e., the rate of CTL expansion.

To assess the effect of compartmentalization on the dynamics, model version (i) is compared to two other settings: (A) The same compartmental model, but with high values of *g* and *η*, i.e. assuming that CTL and infected cells move with a relatively fast rate between compartments (but CTL are again only stimulated in the EF compartment). We refer to this as **model version (ii)**. This is still a compartment model, but the extent of compartmentalization is weak due to fast migrations between compartments. (B) A single compartment model, where the rate of target cell production is given by the sum of the cell production rates in the EF and F compartments, such that that total target cell population size remained the same. We refer to this as **model version (iii)**. This represents complete lack of a compartmental structure. The equations underlying this model are given in the Supplementary Materials.

For each rate of CTL expansion, c, in model version (i), we adjusted the equivalent parameter in model versions (ii) and (iii) such that the equilibrium number of wild-type infected cells in the absence of mutants was identical across all model versions. In this way, we can study the effect of compartmentalization without varying total virus load at the same time, which could blur the picture because variation in virus load alone can impact the dynamics of mutant invasion.

### Properties of model 1, depending on the rate of CTL expansion, c

The strength of the CTL response is an important parameter, because it varies both across different patients, and within one patient as the disease progresses. We vary the parameter *c* (rate of CTL expansion), although similar results are seen if the rate of CTL-mediated killing, *p*, is varied in this model. Note that the parameter *c* is only varied freely in model version (i), and the corresponding parameter values in model versions (ii) and (iii) were adjusted to keep virus load constant across models. The properties of model version (i) depending on the parameter *c* are shown in Figure 3, purple lines.

**Figure 3.**
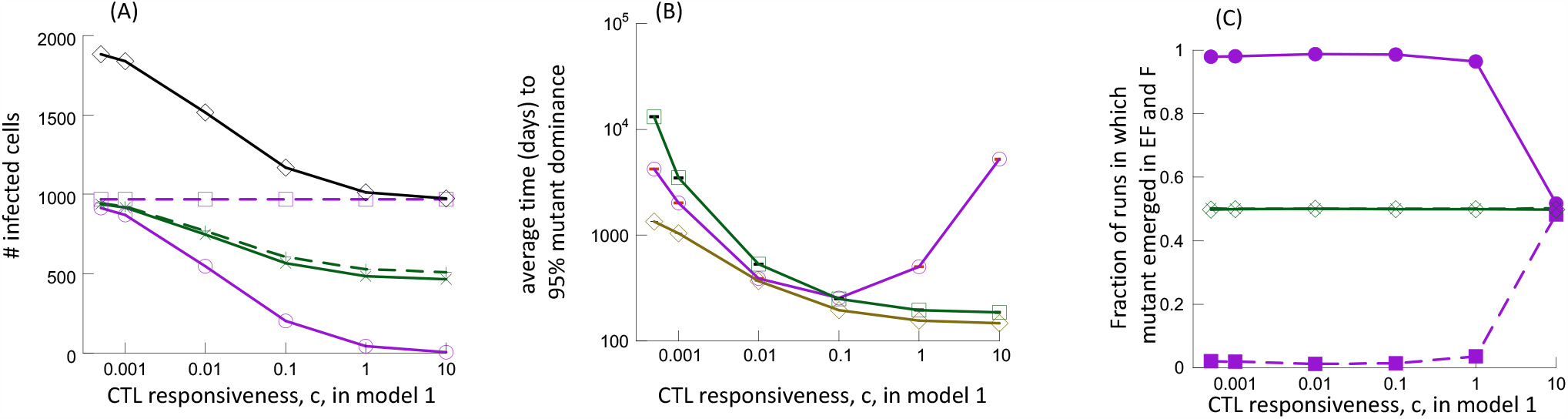
Stochastic evolutionary dynamics of CTL escape, as a function of the rate of CTL expansion or CTL responsiveness, *c*. Gillespie simulations of ODE model (2) were performed. Three model versions are compared, as described in the text. Model version (i) is characterized by strong compartmentalization (*η=g=0.0001*). Model version (ii) still has compartments, but populations move between them with relatively high rates (*η=g=1*). Model version (iii) is a single compartment model. The value of *c* on the x-axis corresponds to model version (i), and is varied from low to high. For each value of c for model (i), we adjust the corresponding value in model versions (ii) and (iii) such that total virus load is identical across the model versions. In this way, we see the effect of compartmentalization only (without additional effects occurring as a result of variation in virus load). (A) Total wild-type virus load (without mutants) as a function of the parameter c in model version (i). These are the values predicted by the ODEs and represent the averages in the stochastic process. The black line denotes total virus load, which is kept identical across all three model versions. The purple and green lines show virus load by compartment for model versions (i) and (ii) respectively. Solid and dashed lines depict the F and EF compartments, respectively. (B) The average time for the CTL escape mutant to reach 95% of the infected cell population in the EF compartment. Purple is model version (i), green model version (ii), and brown model version (iii). Standard errors are shown but are small and difficult to see. (C) Fraction of simulation runs in which the invading CTL escape mutant was generated in the F (dashed line) and the EF (solid line) compartment. Purple is model version (i) and green model version (ii). Parameters were chosen as follows. For the compartmental model version (i): *λ*_*f*_*=500*; *λ*_*e*_*=500*; *δ=0.01*; *a=0.45*; *β*_*e*_*=β*_*f*_*=7×10*^*-5*^; *β*_*1e*_*=β*_*1f*_*=0.99β*_e_; *µ=2×10*^*-5*^; *p=0.05*; *b=1*; *η=0.0001*; *g=0.0001*. For compartmental model version (ii), parameters were identical, but *η*=g=1. For non-compartmental model (iii): *λ=1000*; *δ=0.01*; *a=0.45*; *β7×10*^*-5*^; *β*_*1*_*=0.99β*; *µ=2×10*^*-5*^; *p=0.05*; *b=1*. The initial conditions for the Gillespie simulations were the equilibrium population sizes for the wild-type population, as predicted by the ODEs.

As is typical for virus dynamics models [31], total virus load at equilibrium declines with a faster rate of CTL proliferation, c. However, due to the compartmentalization, this arises almost entirely through a reduction in virus load in the extrafollicular compartment (Figure 3A). Virus load in the follicular compartment is not significantly affected by the rate of CTL proliferation, due to the assumed slow rate at which CTL enter the F compartment.

Figure 3B (purple line) shows that the average time to mutant invasion has a minimum for an intermediate strength of the CTL response, c. This is intuitive and expected. For weak CTL responses, virus load is high in both compartments and the selection pressure is low. While mutants are generated with a relatively fast rate, their ability to rise at the expense of the wild-type virus is limited. For strong CTL responses, there is a high selection pressure in the EF compartment, which facilitates escape mutant invasion. Due to the low virus loads in the EF compartment, however, the probability to generate a mutant is low, which contributes to the increased invasion times. Mutants can be more readily produced in the F compartment, where virus load is higher. However, due to the limited CTL activity in the F compartment, the generated mutants are unlikely to rise to significant levels. They are more likely to go extinct instead, especially because they are assumed to carry an intrinsic fitness cost. For intermediate rates of CTL proliferation, virus load in the EF compartment is intermediate, providing a higher chance to generate the CTL escape mutant, while selection still favors the escape mutant to a sufficient extent. This results in the shortest mutant invasion times.

We determined the contribution of the EF and F compartments to mutant evolution, by tracking the origin of the escape mutants in the computer simulations. When the total mutant infected cell fraction in the EF compartment reached 95%, we determined in which compartment the largest escape clone originated, and thus recorded the fraction of simulation runs in which the largest escape clone originated in the EF and F compartments, respectively. The mutants are largely generated in the EF compartment over a wide range of CTL expansion rates (Figure 3C, purple lines). The reason is that in the F compartment, very few CTL are present and the escape mutant is assumed to carry an intrinsic fitness cost. Hence, the mutant is not likely to sufficiently rise in the follicles and move to the EF compartment. For very strong CTL responses, however, escape mutants originate more frequently in the F compartment. In this parameter regime, the CTL response essentially eliminates virus from the EF compartment, which is only maintained there by influx from the follicles. In this regime, it is thus more likely that escape mutants are produced in the follicle and subsequently move to the extrafollicular compartment than originating in the EF compartment.

### Comparison to control models with less or no compartmentalization

For each value of the parameter c in model version (i), a modified value of c was used in model versions (ii) and (iii) such that total virus load was the same. In this way, we can see how strong compartmentalization alone affects the outcomes, without variation in virus load. In Figure 3, the properties of model version (ii) are shown in green lines, while model version (iii) is shown by a brown line.

Model version (ii) still has compartmental structure, but the fast migration rates between compartments renders the system more mixed, resulting in the virus being almost equally abundant in both the F and the EF locations (Figure 3A, green lines). Hence, both compartments also contribute equally to escape mutant production (Figure 3C, green lines). Comparing the average time to escape mutant dominance in model versions (i) and (ii), different patterns are observed depending on the strength of the CTL response. For relatively strong CTL responses (value of *c* above a threshold), the average time to escape mutant dominance is significantly longer for the strong compartmentalization (model i) than for mixing between compartments (model ii), compare green and purple lines. In other words, strong compartmentalization significantly delays the rate at which CTL escape mutants invade in the EF compartment. The reason is that with strong compartmentalization, the low selection pressure in the F compartment makes it unlikely for mutants to rise there and take over the EF compartment; at the same time, the strong CTL-mediated virus control in the EF compartment results in only infrequent mutant production in that location. In contrast, for weak compartmentalization (model version ii), there is significant selection pressure in both compartments, and virus load in each compartment is intermediate (Figure 3A). This leads to overall faster mutant production, coupled by relatively strong selection.

The situation, however, reverses for weak CTL responses (smaller values of *c*). Now strong compartmentalization (model version i) accelerates mutant invasion in the EF compartment, compared to model version (ii) (Figure 3B). The reason is that in this setting, virus load in the EF compartment is relatively high even for strong compartmentalization, due to the weaker CTL response. This results in more frequent mutant production, and concomitant selection of the produced mutants. For compartmental mixing (model ii), on the other hand, the CTL that are stimulated in the EF compartment can readily move to the F compartment. This makes the total number of CTL in the EF compartment lower, leading to reduced selection pressure and slower mutant invasion.

We now compare the strongly compartmentalized model (i) to the single-compartment model (iii), which lacks any compartmental structure and is more typical of previous mathematical models that considered the evolution of immune escape (Figure 3B, brown line). Now, the rate of escape mutant invasion is fastest (compared to both model versions i and ii). In the strongly compartmentalized model, overall virus load is given by the sum of the low number of infected cells in the EF compartment and the large number of infected cells in the F compartment, due to the difference in CTL activity. To achieve a total virus load that is comparable in the single compartment model (version iii), we have to assume a CTL response of intermediate strength. This results in intermediate virus load in the single compartment, together with significant selection pressure, and this promotes escape mutant invasion. Therefore, given a certain amount of overall virus load, compartmentalized models (in fact both models i and ii) predict a significantly longer time until escape mutant invasion compared to the single compartment model (iii). This again highlights the role of compartmentalization in the delay of CTL escape mutant evolution.

## Discussion and Conclusion

Mathematical models have made major contributions to our understanding of HIV dynamics [32]. Most of this was possible with simplified models that treated the HIV population within an individual as a single, well-mixed system. In some studies, spatial models were used to explore virus spread under more complex assumptions[25,37]. The explicit consideration of virus replication in extra-follicular and follicular compartments, however, has been largely missing. At the same time, recent data indicate that this compartmentalization might be an important factor for determining the extent to which CTL can control the infection. The follicular compartment represents an immune-privileged site in the context of CTL, where the infected cells are shielded from CTL-mediated activity. Failure of CTL to accumulate in the follicles in large numbers has been observed in people living with HIV[5]. In SIV-infected macaques, the presence of strong CTL responses has been shown to result in an unequal distribution of the virus across the two compartment types, with high virus loads in the follicles and low virus loads in the extrafollicular compartments, controlled by CTL [4]. In contrast, the virus population is distributed more equally across compartments in animals with weak CTL responses or in the acute phase of the infection before CTL responses have emerged [4]. These observations have been replicated in a mathematical modeling study, which investigated compartmental dynamics under different assumptions [29].

Here, we have built on this modeling work and have shown that the compartmental structure of the secondary lymphoid tissue has a significant impact on the in vivo dynamics of HIV that can explain seemingly counter-intuitive results. First, the models can account for conflicting observations that demonstrate the importance of CTL for limiting virus load, and at the same time show a lack of correlation between the strength of the CTL response and the estimated life-span of infected cells in HIV-infected patients [28] or SIV-infected macaques [29]. According to the model, the virus decline observed in the blood represents mostly the decline of infected cells in the follicular compartments, because this population is not significantly suppressed by CTL and is therefore dominant, even though CTL-mediated lysis is assumed to suppress virus load in the extrafollicular compartment, limiting the total amount of virus in the host. It would be instructive to measure the rate of virus decline during antiviral therapy in the presence and absence of CTL separately for virus in the extrafollicular and the follicular compartments. According to our theory, CTL depletion should significantly affect the rate of virus decline in the extrafollicular, but not in the follicular compartments.

Second, in the presence of relatively strong CTL responses, the compartmental models predict significantly longer times for the emergence of CTL escape mutants compared to models in which cells mix more readily between the compartments or compared to models without compartment structure. The reason is that in the extrafollicular compartments, where CTL-mediated selection pressure is strong, the chances to generate an escape mutant are small due to low virus load. In the follicular compartment, where virus load is relatively high, escape mutants can be readily generated, but they are unlikely to invade given the lack of a selective advantage in this location. Hence, this results in longer emergence times compared to a model without compartmental structure but same total virus load. This adds to the explanations for the observed slow emergence of CTL escape in untreated people living with chronic HIV, reviewed in the Introduction section. It is interesting to note that the compartmental structure leads to the longest delay in CTL escape mutant emergence compared to mixed models for relatively strong CTL responses. This is in contrast to the notion that unexpectedly long times until escape mutant emergence are indicative of a relatively weak CTL response or predominantly non-lytic CTL activity [19,25]. Moreover, our compartment model is consistent with the observation that CTL escape mutants in people living with HIV arise faster during the acute phase of the infection, in contrast to the slow dynamics during chronic infection [16]. During acute SIV infection, it has been observed that the distribution of virus across the follicular and extrafollicular compartments is more even than during chronic infection, due to less pronounced CTL activity [4]. Hence, the compartmental dynamics that result in delayed CTL escape mutant emergence in our model apply less during acute compared to chronic infection, which could explain the difference in the rate of escape mutant evolution. Thus, our model can reconcile the relatively long emergence times for CTL escape mutants in chronic infection with the presence of strong lytic CTL responses that can limit overall virus load, and simultaneously predicts the observed faster emergence of CTL escape in acute infection.

An important extension of the mathematical work reported here is the broader analysis of viral evolutionary dynamics in the presence of compartmentalization. While there is a clear difference in viral fitness between the two compartment types for CTL escape mutants, the relative fitness of mutants in the follicular and extrafollicular compartments is likely the same for other viral variants that do not contribute to CTL escape. Previous evolutionary work, however, has shown that variation in population sizes across demes can itself change mutant invasion dynamics [38]. Moreover, differences in the stochastic fluctuations in population sizes between the follicular and extrafollicular compartments, which can be brought about by the variation in CTL-mediated activity, can further influence the evolutionary dynamics of the virus [39]. These effects can be quantified by an extension of the mathematical framework presented here.

The formulation of the CTL dynamics in our model is characterized by some uncertainty. The CTL expansion term is phenomenological, simply assuming that the virus-specific CTL population expands at a rate proportional to the amount of their antigen present, *cY*. Other models have assumed true proliferation of the CTL population in response to antigen, where the expansion term is given by *cYZ* [31]. This, however, leads to less stable dynamics with strong oscillations that are not seen in CTL responses *in vivo*. Other models included a certain number of pre-programmed CTL divisions following antigenic stimulation [40,41]. Based on the comparison of the properties of these models, we do not expect that such differences would lead to qualitatively different results in the context of our study. Another assumption we made in our model was that CTL can only become stimulated by antigen in the EF compartments, and not in the F compartments. This is based on the notion that CTL home to the EF compartments and limit their presence in the F compartments [5]. In the context of engineered CTL that can home to the F compartment [42], some degree of antigenic stimulation was observed in the follicles. We do not expect the results to change if the rate of CTL stimulation in the F compartment is relatively low, and the migration of CTL from the F to the EF compartment is relatively large. If, however, significant CTL expansion and activity was assumed to occur in the F compartments, then the model properties would likely become different because this would alter and reduce the differences in virus load across the two compartments. Finally, we assumed that infected CD4 T cells can migrate between the two compartments, going both directions with equal rates. It might be more realistic to assume that CD4 T cell migration occurs predominantly from the F to the EF compartments. This, however, would be unlikely to change conclusions, because the migration of infected CD4 T cells from the F to the EF compartments is key for mutant invasion if the mutant originates in the follicles. Migration in the other directions has no impact on mutant invasion dynamics.

In conclusion, our models suggest that the compartmentalization of virus replication in the secondary lymphoid tissues, with relatively strong CTL-mediated virus control in the extrafollicular compartment, coupled with high virus loads and limited CTL activity in the follicles, can fundamentally influence the dynamics of HIV infection. It can explain the lack of variation in the overall rate of virus decline during antiviral therapy in the peripheral blood, even if CTL-mediated lysis contributes significantly to virus control and if patients differ in the degree of CTL-mediated lytic activity [42]. It can further provide an explanation for the unexpectedly long CTL escape emergence times that have been observed in chronic HIV infection [16-20], even under the assumption that CTL-mediated lysis is a significant contributor to virus control. Therefore, it might be important to investigate more closely the heterogeneity in virus dynamics between the follicular and extrafollicular compartments when studying the dynamics and evolution of HIV *in vivo*, rather than to solely concentrate on plasma virus loads measured in patients.

## Supporting information

Supplementary Materials: Single-Compartment Models

## Data Availability

This paper is based on mathematical models and does not contain novel data.

