## Supplementary Materials: Single-Compartment Models for "Human immunodeficiency virus (HIV) dynamics in secondary lymphoid tissues and the evolution of cytotoxic T lymphocyte (CTL) escape mutants"

The main text defines compartmental models of virus dynamics to study the evolution of cytotoxic T lymphocyte (CTL) escape mutants by human immunodeficiency virus (HIV). In particular, we distinguish between the follicular and extrafollicular compartments in the secondary lymphoid tissues. The results obtained for these models are compared to those seen in a corresponding single-compartment model, which is outlined as follows.

We denote the populations of uninfected cells, infected cells, and CTL by *x*, *y*, and *z*, respectively. The time evolution of these populations is given by the following set of ordinary differential equations:

$$\frac{dX}{dT} = \lambda - dX - \beta XY,$$

$$\frac{dY}{dT} = \beta XY - aY - pYZ,$$

$$\frac{dZ}{dt} = cY - bZ.$$

Similar models have been widely used in the literature, e.g. [1-4]. Target cells are produced with a rate $\lambda$, die with a rate $dX$, and become infected by virus with a rate $\beta XY$ (assuming that virus is in a quasi-steady state). Infected cells are characterized by a basic death rate $aY$, and are killed further by CTL with a rate $pYZ$. The CTL population expands with a rate $cY$ and dies with a rate $bZ$.

In the absence of the CTL, the virus population establishes a persistent infection if its basic reproductive ratio $R_0=\beta\lambda/da>1$. The system then converges to a stable equilibrium, given by $X^{(1)}=a/\beta, Y^{(1)}=\lambda/a - d/\beta, Z^{(1)}=0$. If $c>0$, the CTL response expands in the presence of the infection. In this case, the populations converge towards the following stable equilibrium.

$$X^{(2)} = \frac{a\,b\,\beta - c\,d\,p + \sqrt{(a\,b\,\beta - c\,d\,p)^2 + 4b\beta^2\lambda cp}}{2b\beta^2},$$

$$Y^{(2)} = \frac{b\left(\beta X^{(2)} - a\right)}{cp},$$

$$Z^{(2)} = \frac{\beta X^{(2)} - a}{p}.$$

Next, we incorporate a CTL escape virus strain into the model, described by the subscript "1"; the infected cells are hence denoted by $Y_1$. The modified ODEs are given as follows:

$$\frac{dX}{dT} = \lambda - dX - \beta XY - \beta_1 XY_1,$$

$$\frac{dY}{dT} = \beta XY(1-\mu) - aY - pYZ,$$

$$\frac{dY_1}{dT} = \mu\beta XY + \beta_1 XY_1 - aY_1,$$

$$\frac{dZ}{dt} = cY - bZ.$$

The CTL escape virus strain is characterized by its specific rate of infection, $\beta_1$, and is not killed by the CTL population. It is produced during infection events with a probability $\mu$. We assume that the $R_0$ of both virus strains is greater than unity, and that the CTL population expands. In this case, two outcomes are possible. If $\beta 1>\beta(1-\mu)$, the escape mutant will fixate (with the wild-type virus going extinct). Because the wild-type virus goes extinct and the escape mutant does not stimulate the CTL response, the CTL population also goes extinct in this model. The equilibrium is hence given by $X^{(3)}=a/\beta_1$, $Y^{(3)}=0$, $y1^{(3)}=\lambda/a - d/\beta_1$, $Z^{(3)}=0$. Otherwise, the escape mutant coexists with the wild-type virus and the CTL response against the wild-type virus also persists, i.e. all population sizes are greater than zero. The equilibrium expressions for this outcome are given by the solution of a third degree polynomial and are thus not specified here.


[1] Nowak, M.A. & May, R.M. 2000 *Virus dynamics. Mathematical principles of immunology and virology.*, Oxford University Press.

[2] Perelson, A.S. 2002 Modelling viral and immune system dynamics. *Nature Rev Immunol* **2**, 28-36.

[3] Perelson, A.S. & Ribeiro, R.M. 2013 Modeling the within-host dynamics of HIV infection. *BMC Biol* **11**, 96. (doi:1741-7007-11-96 [pii]
10.1186/1741-7007-11-96).

[4] Wodarz, D., Christensen, J.P. & Thomsen, A.R. 2002 The importance of lytic and nonlytic immune responses in viral infections. *Trends Immunol* **23**, 194-200.
